# Calibration standards and sensitivity limits for fluorescence measurements with the Chi.Bio open-source bioreactor platform

**DOI:** 10.64898/2026.06.29.735387

**Authors:** Alessia Sambruna, Giorgio Tallarico, Marco Cosentino Lagomarsino

## Abstract

Automated platforms such as Chi.Bio enable simultaneous monitoring of optical density and fluorescent reporter expression in 20 ml reactor cultures with controllable pump systems. As such, they provide an appealing option for contemporary gene expression quantification, quantitative physiology, and laboratory evolution and ecology experiments. While optical density calibration for this device is well established, no equivalent calibration framework exists for fluorescence, making quantitative comparison with reference instruments unreliable. Here, we characterize Chi.Bio fluorescence capabilities using fluorescent calibration microspheres and fixed GFP-expressing *S. cerevisiae* and *E. coli* cells, compared with orthogonal plate-reader measurements. We show that microsphere fluorescence is detectable and scales linearly with concentration, whereas the GFP signal from both species falls below the device detection limit. Comparison of background-correction strategies indicates that direct subtraction of a non-fluorescent control measured within the same device yields more reliable fluorescence estimates than the commonly used on-line normalization method. Knowledge of these sensitivity boundaries of the device provides practical guidelines for experimental design of future studies.

## 1 Introduction

Automated platforms such as Chi.Bio enable key measurements in situ at low cost with living cells, combining continuous culturing with tunable light sources and spectrometry [1, 2]. For example, quantitative microbial physiology requires precise control of growth conditions together with continuous measurements of biomass and gene expression [3, 4, 5, 6]. Classic and recent studies have largely relied on cultures grown in batch under near-balanced exponential growth, where physiological variables are measured by periodically sampling the culture and performing offline analyses. While this approach has enabled many of the foundational results in bacterial physiology, it is labor-intensive and provides only sparse temporal measurements. Conversely, plate-reader-based workflows offer higher throughput and online measurements, but mostly provide insufficient control of the growth environment and culture conditions. By integrating culture control with continuous online measurements of optical density and fluorescence in a relatively large culture volume, Chi.Bio offers a promising alternative for quantitative physiology experiments. However, while optical density measurements with Chi.Bio are well established, fluorescence measurements have not been systematically validated against reference instruments, limiting their quantitative interpretation.

The Chi.Bio has been extensively used for optical density measurements, as a chemostat and turbidostat, and as an optogenetic tool [7, 8, 9, 10, 11, 12, 13, 14, 15, 16, 17, 18]. Several studies have also leveraged its fluorescence capabilities, primarily in induction experiments where fluorescence is tracked as a relative time-dependent signal [19, 20, 21]. To reduce device-to-device variability, the original publication proposed a ratiometric normalization approach in which the fluorescence signal is divided by the total excitation intensity [2]. However, this approach has not been validated against reference instruments with controlled samples, and its performance for absolute quantification across devices remains uncharacterized.

Here, we systematically characterize the fluorescence measurement capabilities of Chi.Bio using fluorescent calibration microspheres, fixed GFP-expressing *S. cerevisiae* and *E. coli* cells as controlled samples, as well as living *E. coli* cells. We identify the conditions under which reliable fluorescence quantification is achievable, characterize the failure modes of the proposed normalization approach, and define the sensitivity boundaries of the device, providing practical guidelines for users planning fluorescence experiments with this platform.

## 2 Results

### 2.1 Fluorescent microspheres produce reliable, concentration-dependent signals, detectable above background

We first tested Chi.Bio fluorescence detection using fluorescent calibration microspheres as stable, well-characterized reference samples with known excitation and emission properties (Section 3). We used yellow-green and pink fluorescent microspheres and non-fluorescent controls of comparable size and material (Methods). A schematic of the Chi.Bio setup is shown in Figure S1.

Both microsphere types produced signals clearly distinguishable from the non-fluorescent reference, scaling linearly with particle concentration (Figure 1A). The net fluorescence intensity, obtained by subtracting the non-fluorescent control signal, also scaled linearly with concentration with intercepts close to zero, confirming effective background removal (Figure 1B).

**Figure 1:**
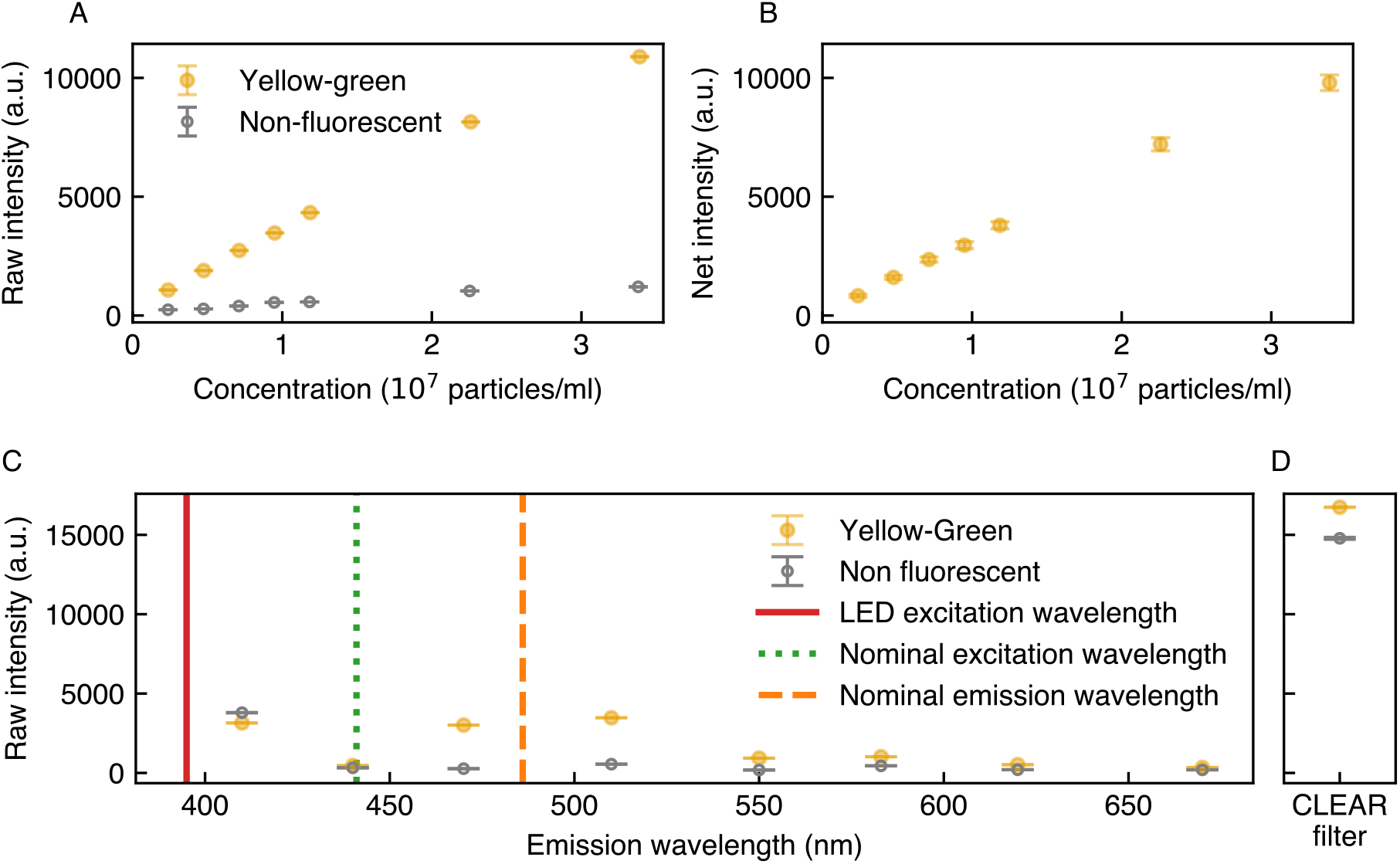
Yellow-green fluorescent microspheres produce reliable concentrationdependent signals, detectable above background. (A) Raw fluorescence intensity measured by the Chi.Bio spectrometer for yellow-green microspheres (yellow filled circles) and non-fluorescent reference beads (grey open circles) as a function of concentration (excitation/emission: 395/510 nm, gain: 512, power: 0.1, device: 0). (B) Net intensity for yellow-green microspheres as a function of concentration. The value was calculated via direct subtraction of the non-fluorescent signal from the fluorescent signal in panel A. (C) Raw fluorescence intensity detected in device 0 using a 395 nm excitation wavelength for yellow-green beads, shown as a function of emission wavelength (left) and using the Clear filter (right). A distinct emission peak is observed around the nominal emission wavelength, separated from the background.

Full emission spectra revealed an intensity peak near the excitation wavelength, attributable to scattered light given the 90° geometry of the detector relative to the LED (Figure 1C). A distinct fluorescence emission peak was additionally visible for yellow-green microspheres around the nominal emission wavelength, whereas the pink microsphere signal was weaker, consistent with their lower brightness (Figure S2). Careful selection of the discrete excitation and emission wavelengths allowed by the device is therefore necessary to avoid spectral overlap with the scattering peak.

These results show that Chi.Bio can reliably detect fluorescence from sufficiently bright samples under optimized parameter settings. However, translating raw signals into quantitative per-particle values requires a normalization approach, which we evaluate in the following section.

### 2.2 The ratiometric normalization based on the Clear channel introduces nonlinearities at low and high concentration regimes

To assess whether the normalization proposed in [2] yields a reliable per-particle fluorescence estimate, we evaluated its behavior across a concentration range of 0.2 *×* 10^7^ to 3.4 *×* 10^7^ particles/ml.

Figure 2A displays the raw intensity measured with the Clear sensor for pink and non-fluorescent microspheres, while Figure 2B shows the normalized intensity for the same samples. The plots show a separation of the fluorescent signal from the background. However, two main problems arise: at low concentrations, background terms dominate introducing non-linearity, while at high concentrations, saturation of the Clear channel at the maximum detectable value (65535) offsets the normalization. This behavior is consistent with the known optical properties of the device, in which excitation light leaks through emission filters to a degree that depends on concentration ^1^. The origin of this non-linearity can be understood analytically by expressing the normalized intensity as

**Figure 2:**
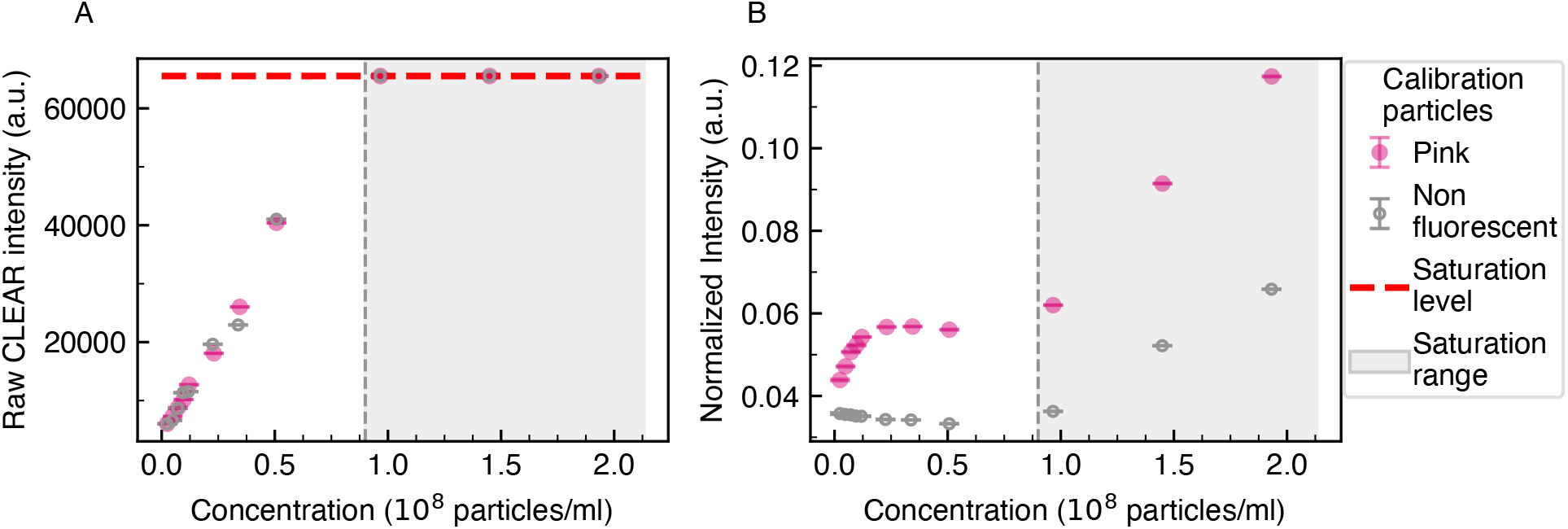
Ratiometric normalization based on the Clear sensor introduces non-linearities at low and high concentration regimes. (A) Raw intensity measured with the Clear channel for pink (pink filled circles) and non-fluorescent (grey empty circles) microspheres. The signal saturates at higher concentrations (grey area) (excitation/emission: 523 nm/CLEAR, gain: 512, power: 0.1, device: 0). (B) Normalized intensity (raw intensity measured within the 620 nm emission channel divided by the Clear channel signal) for pink microspheres. The signal is linear at low-to-moderate concentrations but exhibits saturation effects at higher concentrations as the Clear signal approaches its maximum (65535). All measurements: excitation 523 nm, gain *×*512, power 0.1.

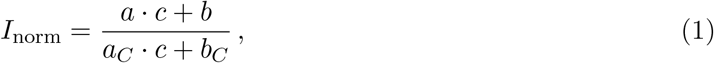

where *c* is the particle concentration. The parameters *a* and *a*_*C*_ represent the concentrationdependent contributions, including fluorescence, reflection, transmission, and scattering effects, for the filtered and unfiltered channels respectively. The intercepts *b* and *b*_*C*_ account for the concentration-independent background in each channel.

Eq. (1) shows that parameters need to be carefully explored for each experiment, and that identifying the optimal concentration range is important to avoid both saturation and low-concentration effects. As shown in the previous section, direct subtraction of a matched non-fluorescent control avoids these problems and appears as a more robust alternative where a suitable control is available. Note however that this comparison must be carried out for samples measured within the same reactor, or carefully calibrated for variability across reactors, posing a practical constraint for many experimental setups involving living cells.

### 2.3 There are limitations in the separation of yeast cells expressing Rpl5-GFP from the background

Having established that Chi.Bio can detect fluorescence from sufficiently bright calibration standards, we next tested whether GFP-expressing biological cells fall within the detectable range of the device. Figure 3A shows fluorescence measurements from fixed yeast cells expressing Rpl5-GFP, a GFP-tagged highly expressed ribosomal protein, at different concentrations, compared with wild-type cells for one single device (device 0) averaged over three biological replicates.

**Figure 3:**
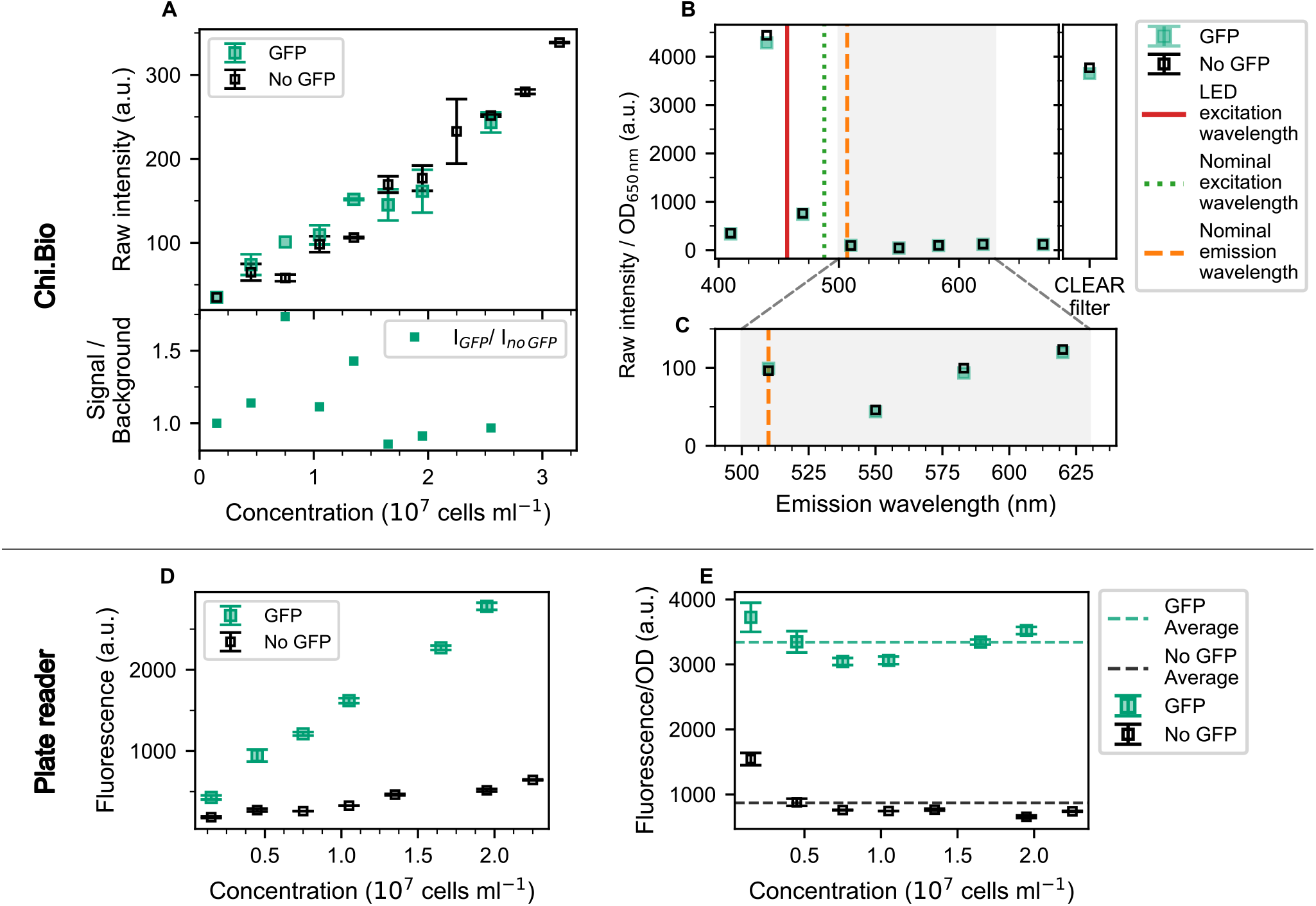
There are limitations in the separation of yeast cells expressing Rpl5-GFP from the background. (A) Top: Raw fluorescence intensity as a function of cell concentration for GFPexpressing yeast cells and non-fluorescent control cells averaged across three biological replicates (Chi.Bio, gain 512, power 0.01, device 0, excitation/emission: 457/510 nm). Error bars: standard deviation across replicates. (A) Bottom: Signal-to-background ratio (Raw I_GFP_/Raw I_no GFP_). (B) Raw fluorescence spectra normalized by OD_650_ for GFP-expressing and wild-type *S. cerevisiae* cells at the highest concentration (≈ 2.5 *×* 10^7^ cells/ml), one biological replicate. (C) Zoom on the emission region shows no discernible GFP peak. Nominal GFP excitation (490 nm) and emission (507 nm) wavelengths are shown as dashed and dotted vertical lines; excitation wavelength (457 nm) as a solid red line. (D) Fluorescence of the same samples as in panel A-C measured in a monochromator-based plate reader (excitation/emission: 457/510 nm) (E) Fluorescence of the same samples as in panel AC measured in a monochromator-based plate reader (excitation/emission: 457/510 nm) normalized by OD_600_.

The fluorescence signal measured from GFP-expressing cells (green symbols) overlapped with that of the wild-type control strain (grey symbols) across the investigated concentration range. Within experimental variability, no systematic separation between the two conditions was observed, indicating that GFP-derived fluorescence was not resolvable above background under these measurement settings (Figure 3A, top).

The signal-to-background ratio (Figure 3A, bottom) was computed as the ratio between the binned mean fluorescence of GFP-expressing cells and that of the wild-type control at matched concentration intervals. Across the investigated range, the ratio remained close to unity, indicating that fluorescence from GFP-expressing cells was not distinguishable from background under these settings.

Inspection of the full emission spectra, following the same approach used for the fluorescent microspheres (Section 2.1), supports this conclusion (Figure 3B). The spectra of GFP-expressing and wild-type cells are nearly identical across all measured wavelengths. Although small differences are observed at several filters, these variations are distributed throughout the spectrum rather than being localized to the expected GFP emission region, suggesting they arise from residual concentration variability or sample preparation differences rather than genuine GFP signal (Figure 3C). Adjustment of gain and excitation power does not overcome the intrinsic sensitivity limit of Chi.Bio (Figure S4). Applying the ratiometric normalization described in the previous section did not improve discrimination between GFP-expressing and wild-type cells.

In contrast, plate reader measurements of the same samples produced a clear, concentrationdependent fluorescence signal well separated from the wild-type background (Figure 3D, E), confirming that GFP expression levels were sufficient for detection by a more sensitive instrument.

### 2.4 GFP-expressing *E. coli* shows marginal signal separation, limited by interdevice variability

Having established that GFP fluorescence from fixed yeast cells falls below the Chi.Bio detection limit, we extended the characterization to fixed GFP-expressing *E. coli* cells, which produce a higher number of GFP molecules per cell during exponential growth [22].

Raw fluorescence intensities scaled linearly with cell concentration but showed no clear separation between GFP-expressing and wild-type cells (Figure 4A). The signal-to-background ratio fluctuated around unity across the investigated concentration range, with only two concentrations exceeding the 10% discrepancy threshold (Figure 4B).

**Figure 4:**
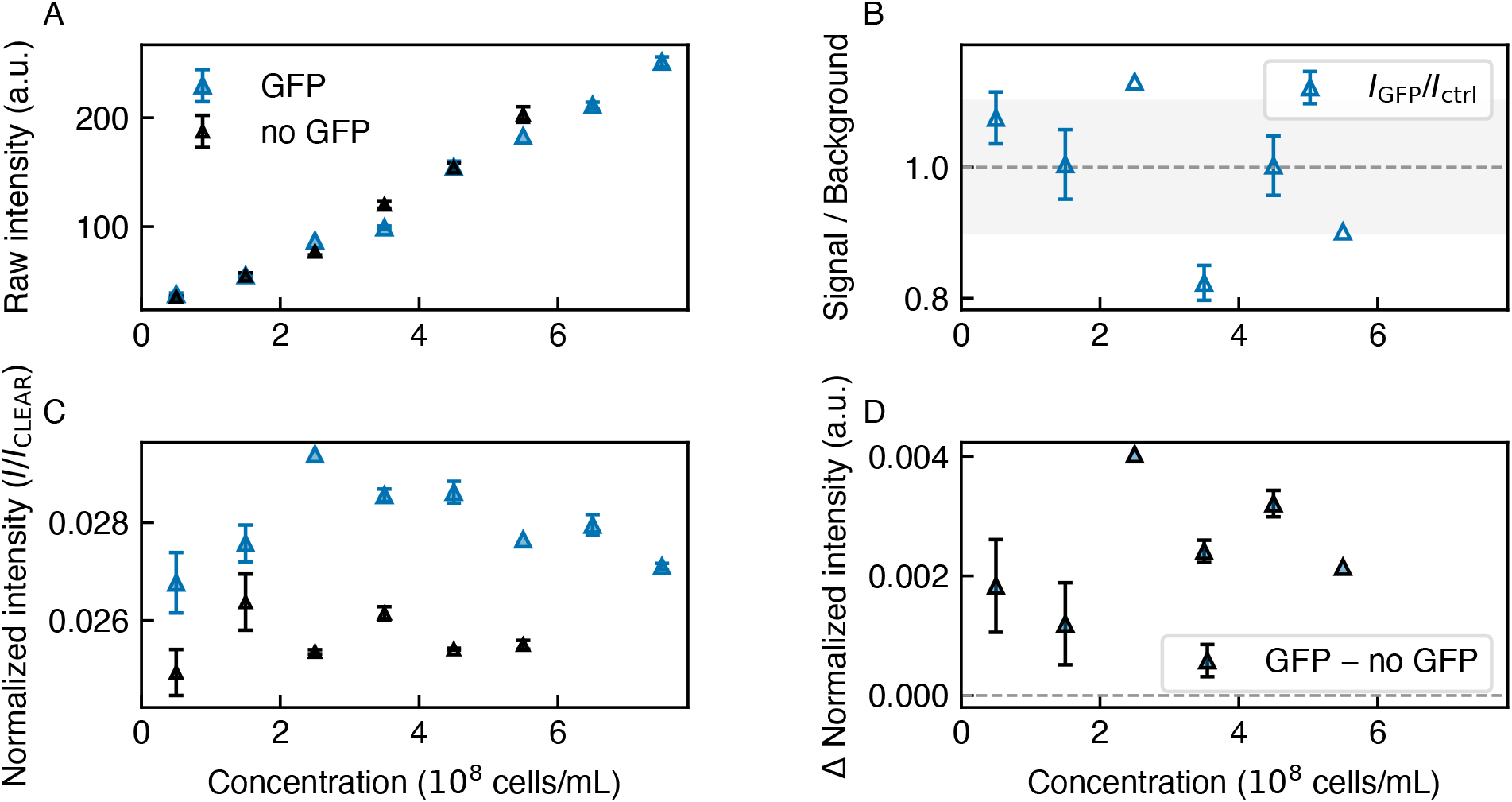
GFP-expressing *E. coli* shows marginal signal separation, limited by interdevice variability. (A) Raw fluorescence intensity as a function of cell concentration for GFP-expressing *E. coli* cells and non-fluorescent control cells averaged across three biological replicates. Error bars: standard error across replicates. (B) Signal-to-background ratio. (C) Fluorescence intensity normalized by dividing by the Clear signal as a function of cell concentration. Error bars: standard error across replicates. (device 0, gain: *×* 512, power: 0.01) (D) Net normalized intensity (panel C, GFP minus wild-type).

After ratiometric normalization, a marginal separation between GFP-expressing and wild-type signals became apparent (Figure 4C, D). However, the magnitude of this separation was comparable to the inter-device variability, as quantified in the following section, limiting its reliability for quantitative interpretation.

### 2.5 Inter-device variability limits reliable fluorescence quantification across devices

To quantify the impact of device variability on the background-subtracted signal, we computed the ratio between the net fluorescence signal and the inter-device variability *σ*_device_, defined as the standard deviation across devices (Figure 5B). The ratio fluctuates around a mean value of 3.3, indicating that the net signal exceeds the inter-device noise on average, but does not remain consistently above a reliable threshold across concentrations, affecting robust quantification.

**Figure 5:**
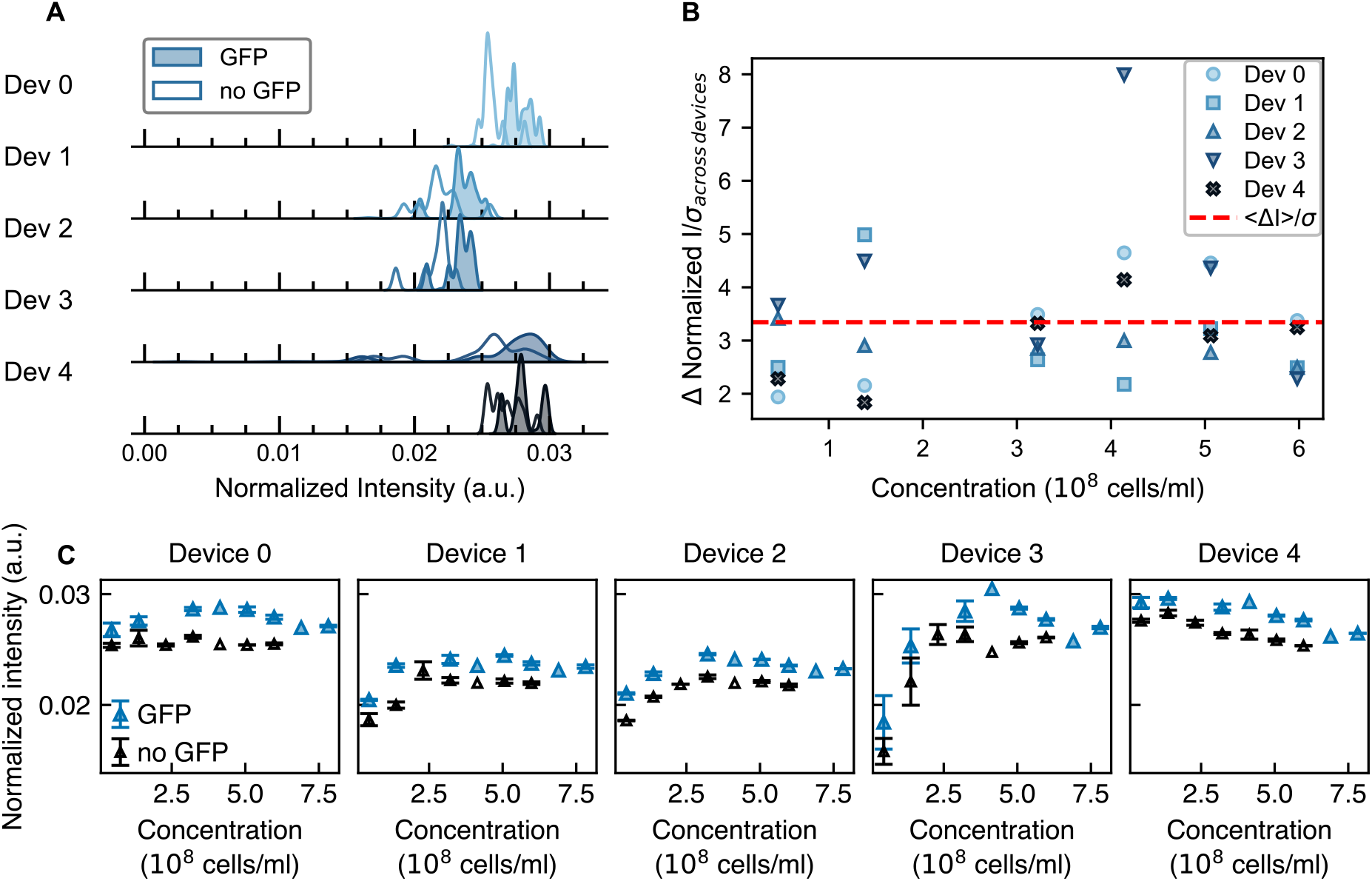
Inter-device variability limits the effective separation of bacterial fluorescence signal from background. (A) Kernel Density Estimation (KDE) of the normalized intensity for GFP-expressing cells (filled) and wild-type background (empty), shown for each device. (B) Net intensity (GFP minus wild-type, normalized) divided by the global inter-device variability (*σ*_device_). (C) Normalized fluorescence intensity as a function of cell concentration for GFP-expressing and wild-type strains, shown separately for each device. Data: mean across three biological replicates; error bars: SEM (power: 0.01, gain: *×*512).

To further characterize this variability, we examined the distribution of normalized fluorescence intensity across devices using kernel density estimation (KDE). In the absence of noise, one would expect a narrow distribution around a single value representing fluorescence per cell. Instead, the distributions of GFP-expressing and wild-type cells are separated from each other, but their peak positions shift across devices, reflecting hardware-specific offsets that persist even after normalization (Figure 5A). Figure 5C reveals non-trivial device-specific concentration trends that cannot be corrected by a simple additive offset, further complicating cross-device comparisons.

In contrast, calibration microspheres show a strikingly different behavior: estimated distributions for fluorescent and non-fluorescent microspheres remain clearly separated across all devices despite hardware-specific offsets, confirming that the device variability observed for bacterial samples reflects genuine sensitivity limitations rather than instrumental failure (Figure S6).

These results indicate that per-device calibration using a fluorescent reference standard measured under identical conditions is required before cross-device comparisons can be made reliably.

In contrast, plate reader measurements of the same samples produced a clear, concentrationdependent fluorescence signal well separated from the wild-type background (Figure S5), confirming that GFP expression levels were sufficient for detection by a more sensitive instrument.

## 3 Discussion

Our results provide a systematic characterization of Chi.Bio fluorescence measurement capabilities and define the conditions under which reliable quantification is achievable. Fluorescent calibration microspheres produced linear signals at the expected excitation and emission ranges, showing that the device can detect fluorescence from sufficiently bright samples. In contrast, fixed GFP-expressing *S. cerevisiae* cells produced no detectable signal above background, and *E. coli* showed only marginal separation.

The primary cause of the limited sensitivity for biological samples is the optical architecture of the device. Broad-spectrum LEDs leak excitation light through the emission filters, producing a concentration-dependent background that is difficult to separate from the fluorescence signal, particularly when excitation and emission wavelengths are close [1]. The 90° geometry between the LED and the spectrometer further increases the contribution of scattered light to the detected signal. These limitations are intrinsic to the current hardware design and confirmed by the platform developers ^2^. Importantly, plate reader measurements of the same samples produced clear, concentration-dependent signals, confirming that insufficient GFP expression is not the explanation.

The ratiometric normalization proposed in [2] can introduce non-linearities at low and high concentration regimes and does not adequately correct for inter-device offsets. This failure is particularly problematic in dynamic experiments with periodic dilutions, where the normalization generates artifactual signal changes that can be mistaken for biological responses. Direct subtraction of a non-fluorescent control measured within the same device is a more robust alternative.

Based on our results and direct correspondence with the platform developers, we recommend the following experimental practices for users planning fluorescence measurements with Chi.Bio. Nonfluorescent media should be used whenever possible, as autofluorescence from rich media such as LB significantly increases background. If possible, experiments should be designed so that fluorescence is induced from a zero-signal baseline, starting with uninduced cells and subtracting the pre-induction baseline from all subsequent measurements, rather than comparing two steady-state conditions. Additionally, per-device calibration using a fluorescent reference standard is required before cross-device comparisons can be made. Finally, the expected fluorescence signal should be verified in a more sensitive reference instrument before committing to Chi.Bio measurements.

For applications requiring higher sensitivity, hardware modifications may be considered. Coupling a dedicated visible-range spectrometer to the reactor via fiber optic, combined with a high-pass filter to reduce excitation leakage, could substantially improve the signal-to-noise ratio. Alternatively, some groups have integrated external flow cytometers with Chi.Bio setups to perform sensitive fluorescence measurements while retaining the platform’s culture control capabilities [23, 24]. The open-source nature of Chi.Bio makes both software and hardware modifications accessible in principle to groups with sufficient technical expertise.

## Materials and Methods

### Data and code availability

Source code for measurement acquisition and analysis scripts will be made publicly available on github.

### Chi.Bio platform and data acquisition

Data were acquired using five Chi.Bio reactors connected to a shared microcontroller. Measurements were performed using custom acquisition routines integrated into the original Chi.Bio control software, available at https://github.com/HarrisonSteel/ChiBio. The thermostat and peristaltic pumps were not activated in this work, as all samples were measured at ambient temperature without flow.

### Calibration microspheres

Three types of polystyrene microspheres were used (Table 1): yellow-green fluorescent microspheres (Fluoresbrite YG Carboxylate Microspheres, nominal ex/em: 441/486 nm, diameter 1.00 *±* 0.03 *µ*m; [25]), pink fluorescent microspheres (PS-FluoRed microparticles, nominal ex/em: 530/607 nm, diameter 0.98 *±* 0.03 *µ*m; [26]), and non-fluorescent amino-polystyrene particles (AP-10-10, diameter 1.0–1.4 *µ*m; [27]).

**Table 1:** Calibration microspheres used in this work.

| Type | Designation | Nominal ex/em (nm) |
| --- | --- | --- |
| Yellow-green | Fluoresbrite YG [25] | 441/486 |
| Pink | PS-FluoRed [26] | 530/607 |
| Non-fluorescent | AP-10-10 [27] | — |

#### Sample preparation

Stock suspensions were diluted in Milli-Q water. For yellow-green microspheres, 50 *µ*l of stock was added to 1 ml of Milli-Q water; for non-fluorescent microspheres, 27 *µ*l was used to match particle concentration. For pink microspheres, which produced a weaker signal at equivalent concentrations, a more concentrated working solution was prepared (0.250 *µ*l stock per 1 ml), with non-fluorescent microspheres matched accordingly (127 *µ*l per 1 ml). Aliquots of varying volumes were transferred into Chi.Bio-compatible glass vials prefilled with 20 ml Milli-Q water to generate the final dilution series.

#### Fixed *S. cerevisiae* cells

Two strains were used (Table 2): wild-type BY4741 and BY4741-RPL5-GFP, in which GFP (S65T) is fused to the ribosomal protein Rpl5 [28]. Cells were grown from glycerol stock on YPD agar, and a single colony was used to inoculate a 10 ml overnight pre-culture in YPD. The saturated pre-culture was diluted into 200 ml fresh YPD in a 1 L flask and grown at 30°C with shaking until exponential phase (OD_600_ ≈ 0.3). Cultures were divided into 50 ml Falcon tubes of varying volumes to generate samples across a range of cell concentrations. Fixation was performed as follows: centrifugation at 4000 rpm for 6 min, resuspension in 20 ml 4% (w/v) paraformaldehyde (PFA) in PBS, incubation at room temperature with gentle shaking for 1 h, and final resuspension in 20 ml PBS after a second centrifugation step. Fixed suspensions were transferred into sterilized Chi.Bio-compatible vials. Three biologically independent replicates were performed; specific OD values at harvest are provided in Table 3.

**Table 2:** Biological materials used in this work.

| Reagent type | Designation | Additional information | Source |
| --- | --- | --- | --- |
| Strain (*S. cerevisiae*) | BY4741 | Non-fluorescent background | SDG |
| Strain (*S. cerevisiae*) | BY4741-RPL5-GFP | Expresses GFP (S65T) | [28] |
| Strain (*E. coli*) | TOP10 (carrying pAG306-pTet07-nLuc plasmid) | Non-fluorescent background | [31] |
| Strain (*E. coli*) | TB204 | Expresses sfGFP | [29] |
| Strain (*E. coli*) | BW25113 | Wild-type background (shifts) | [32] |
| Strain (*E. coli*) | P1L | rrnBP1-GFP reporter (shifts) | [33] |

**Table 3:** Starting OD_600_ values at harvest for each biological replicate (*S. cerevisiae*).

| Replicate | Strain | OD<sub>600</sub> at harvest | Volumes prepared (ml) |
| --- | --- | --- | --- |
| 1 | BY4741 | 0.415 | 10, 20, 30, 35, 40, 50 |
| 1 | BY4741-RPL5-GFP | 0.330 | 10, 20, 30, 35, 40, 50 |
| 2 | BY4741 | 0.755 | 3, 5×2, 10×2, 15×2, 17, 20×2, 25 |
| 2 | BY4741-RPL5-GFP | 0.694 | 3, 5×2, 10×2, 15×2, 17, 20×2, 25 |
| 3 | BY4741 | 0.318 | 5, 10, 15, 20, 30, 40, 50 |
| 3 | BY4741-RPL5-GFP | 0.253 | 5, 10, 15, 20, 30, 40, 50 |

**Table 4:** Starting OD_600_ values at harvest for each biological replicate (fixed *E. coli*).

| Replicate | Strain | OD<sub>600</sub> at harvest | Volumes prepared (ml) |
| --- | --- | --- | --- |
| 1 | no GFP | 0.271 | 5, 10, 20, 30, 40, 50 |
| 1 | GFP | 0.403 | 5, 10, 20, 30, 40, 50 |
| 2 | no GFP | 0.299 | 5, 10, 20, 30, 40, 50 |
| 2 | GFP | 0.389 | 5, 10, 20, 30, 40, 50 |
| 3 | no GFP | 0.255 | 5, 10, 20, 30, 40, 50 |
| 3 | GFP | 0.349 | 5, 10, 20, 30, 40, 50 |

#### Fixed *E. coli* cells

Two strains were used (Table 2): wild-type TOP10 (carrying pAG306pTet07-nLuc plasmid) and TB204 expressing sfGFP [29]. Cells were grown from glycerol stock in LB medium overnight at 30°C with shaking, then diluted into 200 ml fresh LB and grown to OD_600_ ≈ 0.3. Sample preparation and fixation followed the same protocol as for yeast, with the addition of a PBS wash step before PFA resuspension. Three biologically independent replicates were performed.

#### Reference measurements

##### Plate reader

Fluorescence measurements were performed using a monochromator-based Tecan Infinite® 200 PRO multimode plate reader. Samples of 150 *µ*l were loaded into 96-well plates in triplicate. Excitation and emission wavelengths were matched to Chi.Bio settings: 395/510 nm for yellow-green microspheres and fixed cells, 523/620 nm for pink microspheres. For fixed cell experiments, measurements were repeated at multiple gain values; gain settings are reported in the corresponding figure legends.

#### Reference spectrophotometer

Optical density at 600 nm was measured using an Ultrospec 2100 Pro spectrophotometer (Biochrom) in a 1 cm path-length cuvette. Blanks were prepared using the appropriate medium for each sample type.

### Data analysis

#### Concentration binning

Because cell concentrations did not assume identical values across biological replicates, fluorescence measurements were discretized into uniform OD intervals of ΔOD = 0.1 to enable consistent comparison. This bin width reflects the resolution of the experimental dilution scheme. Cell number was estimated using the conversion 1 OD_600_ ≈ 3 *×* 10^7^ cells/ml for *S. cerevisiae* [30].

#### Net fluorescence estimation

Within each bin, mean fluorescence (*Ī*), standard deviation (*σ*), and standard error of the mean 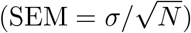 were computed independently for GFP-expressing and control samples. The net signal was calculated as Δ*I* = *Ī*_GFP_ − *Ī*_CTRL_, with propagated uncertainty:

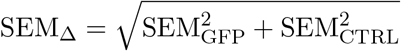

## Acknowledgments

We thank the members of Marco Cosentino Lagomarsino laboratory for valuable discussions and feedback. We are also grateful to Harrison Steel, Luca Ciandrini and his group, Gabriele Micali and his group, Ashley Nord, Pietro Cicuta, and Bianca Sclavi for valuable discussions and suggestions.

We thank Francesco Mantegazza and Domenico Salerno (Università degli Studi di Milano-Bicocca) for providing yellow-green fluorescent and non-fluorescent calibration microspheres, and Fabio Giavazzi (Università degli Studi di Milano) for providing fluorescent pink calibration microspheres.

Bacterial strain TB204 was kindly provided by the Gabriele Micali laboratory.

## Supplementary figures

**Figure S1:**
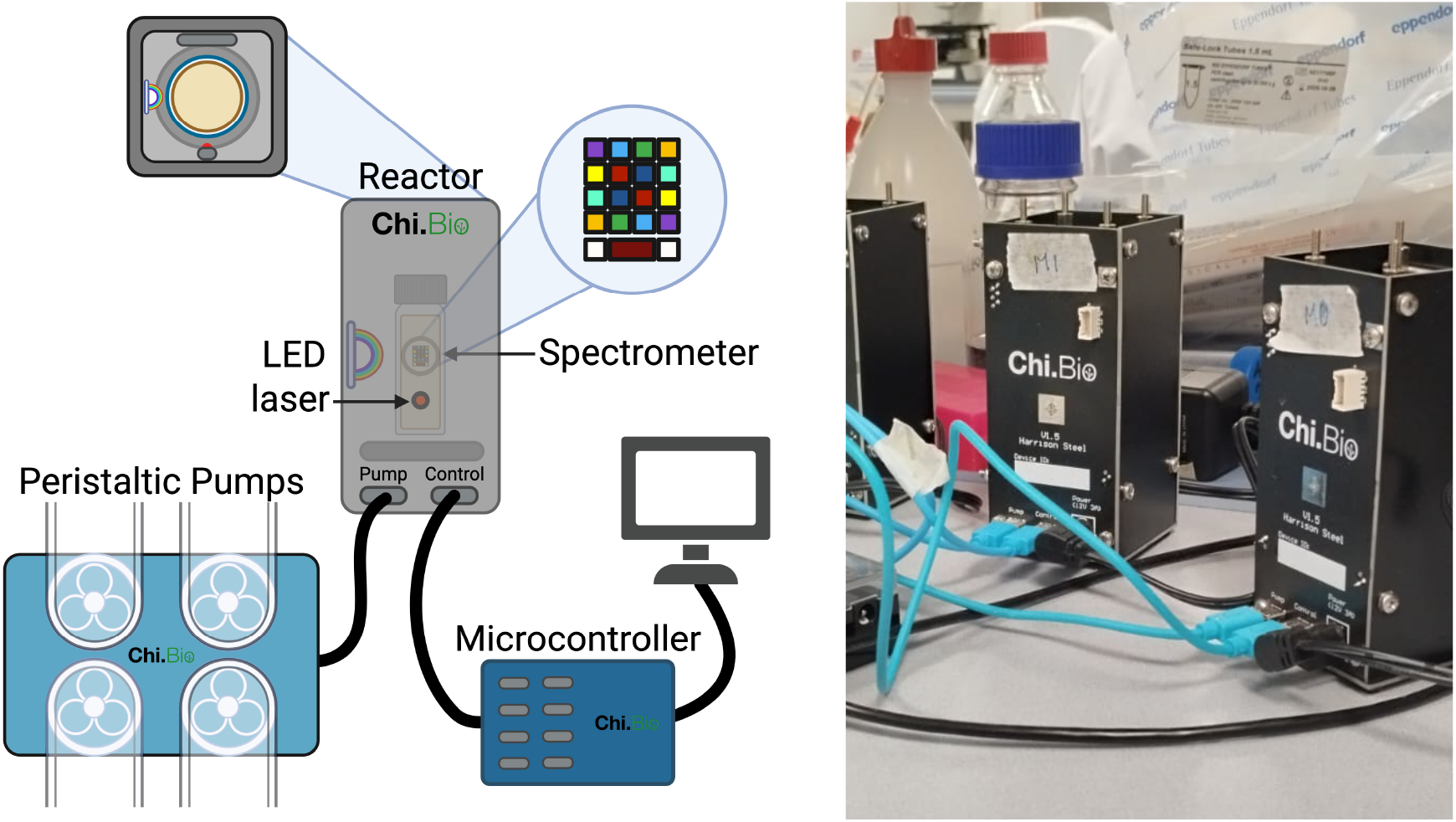
The Chi.Bio device enables automated measurement and monitoring of biological samples. (*Left*) Scheme of the Chi.Bio setup. Each reactor is connected to a peristaltic pump board and to a shared microcontroller, which interfaces with a computer for experimental control. Each reactor contains a vial holding the sample and is equipped with an LED, a laser, and a spectrometer. The figure was created using BioRender, based on the descriptions and figures from ref. [2] and [34]. (*Right*) Picture of two of the five reactors used in this work.

**Figure S2:**
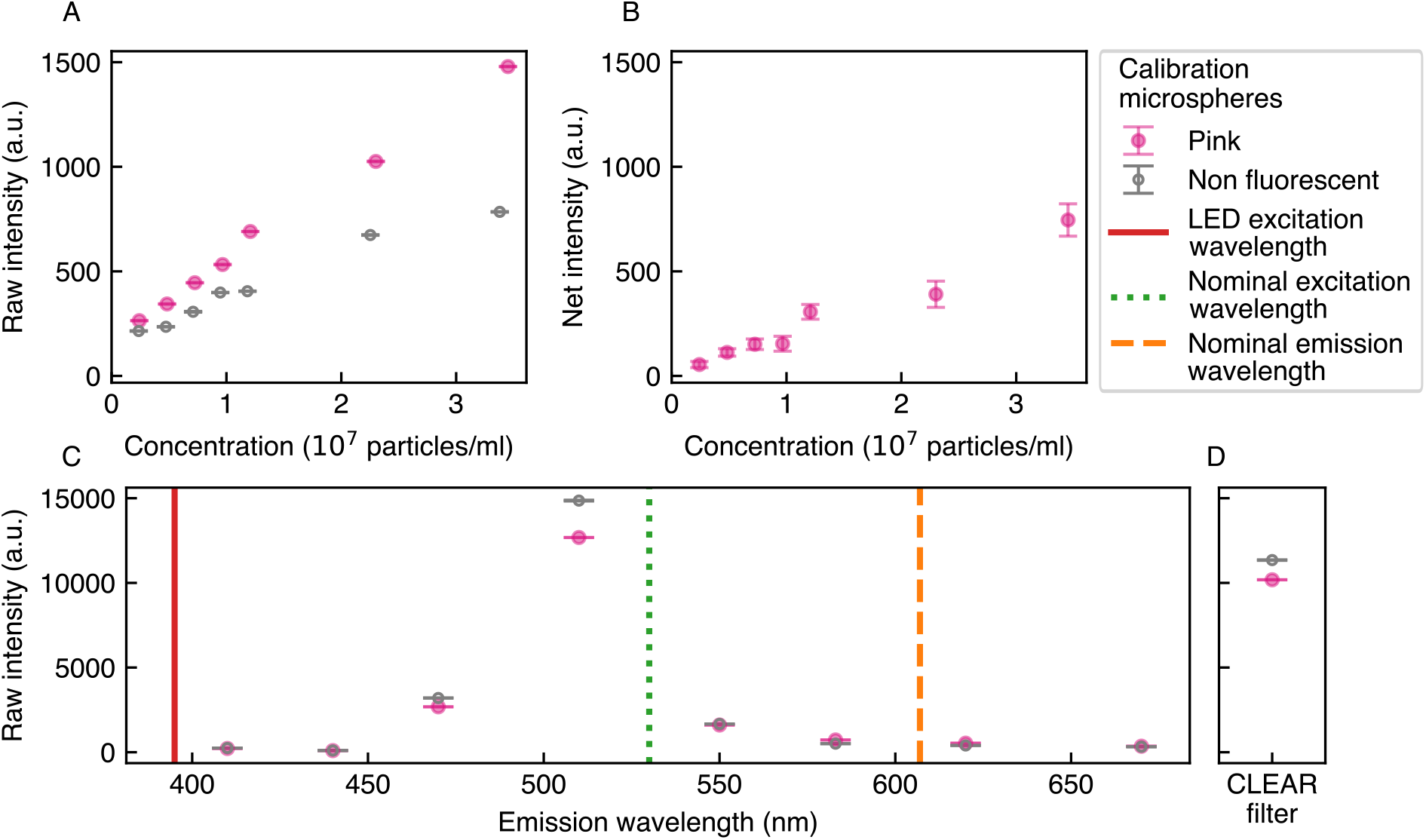
Pink fluorescent microspheres produce reliable, concentration-dependent signals detectable above background. (A) Raw fluorescence intensity measured by the Chi.Bio spectrometer for pink microspheres (pink filled circles) and non-fluorescent reference beads (grey open circles) as a function of concentration (excitation/emission: 523/620 nm, gain: 512, power: 0.1, device: 0). (B) Absolute intensity for pink microspheres as a function of concentration. The linear fit (dashed line) shows that the intercept is approximately zero, indicating negligible background signal. (C) Raw fluorescence intensity detected in device 0 using a 523 nm excitation wavelength for pink beads, shown as a function of emission wavelength (left) and using the Clear filter (right).

**Figure S3:**
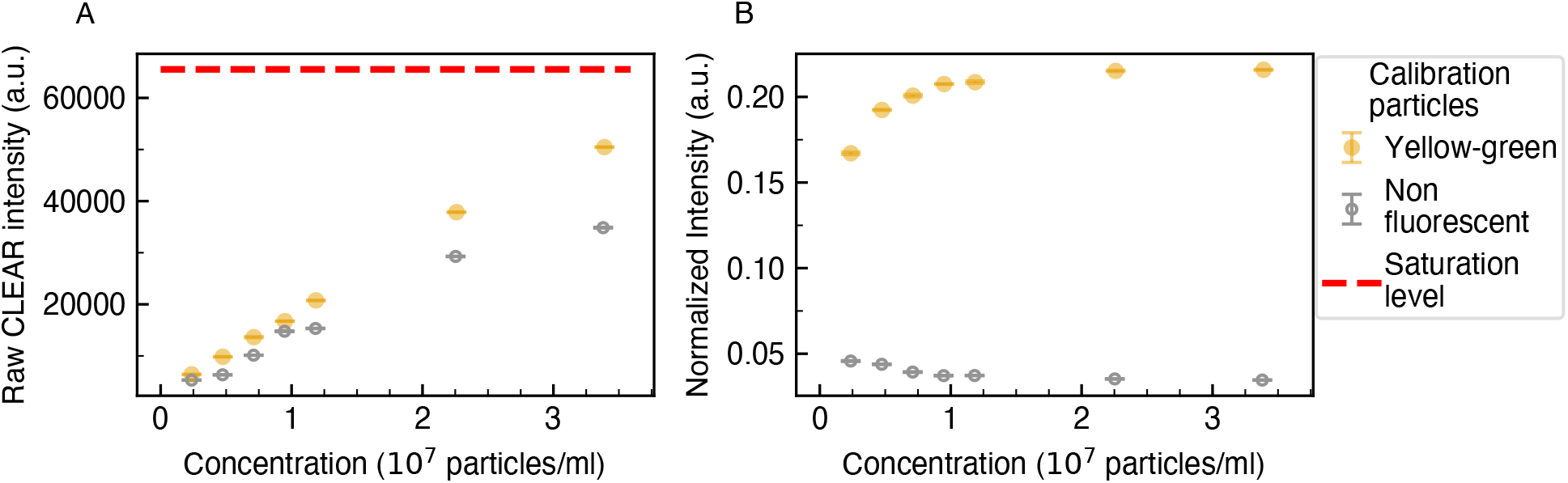
The raw intensity normalized by the Clear signal displays non-linear behavior for yellow-green microspheres. (A) Raw intensity measured with the Clear sensor for yellowgreen (yellow filled circles) and non-fluorescent (grey empty circles) microspheres. (B) Normalized intensity for yellow-green microspheres. All measurements: gain *×*512, power 0.1.

**Figure S4:**
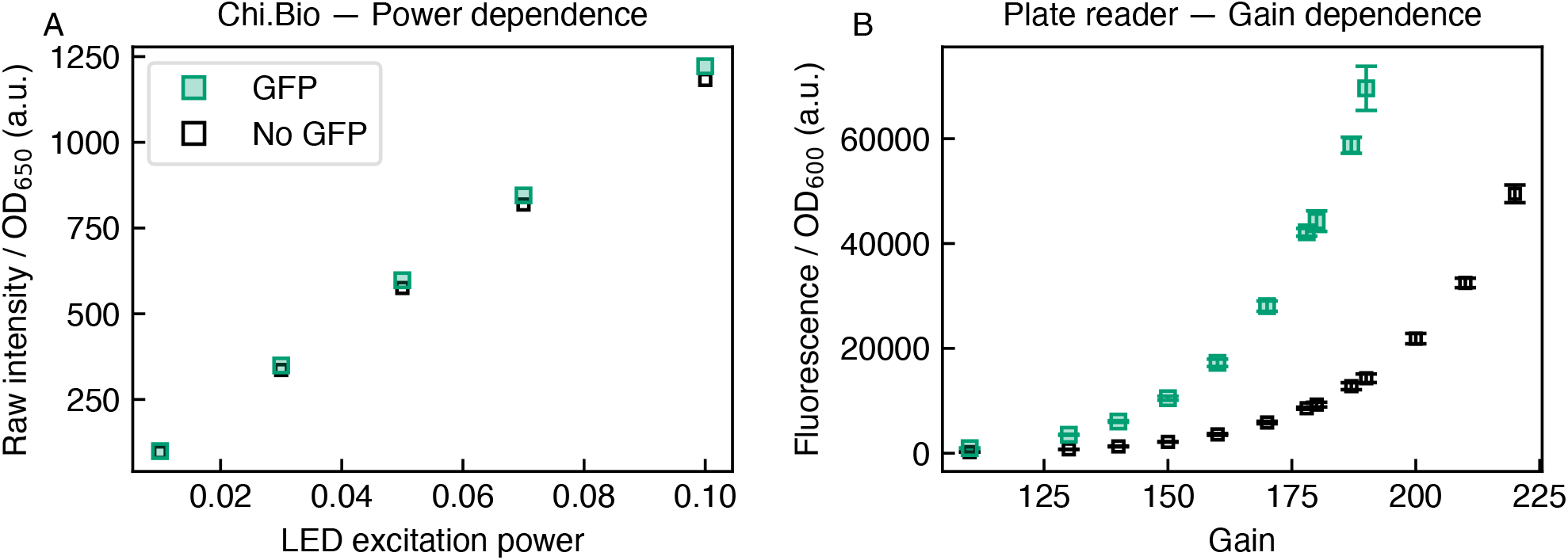
Adjusting gain and excitation power does not improve GFP signal separation in Chi.Bio, in contrast to plate-reader measurements of the same samples. (A) Raw fluorescence normalized by OD_650_ in Chi.Bio (device 0) as a function of LED excitation power for GFP-expressing (green) and wild-type (grey) yeast cells. Signal scales linearly with power but separation remains below inter-device variability. (B) Fluorescence normalized by OD_600_ in a plate reader as a function of gain. Increasing gain improves separation, confirming that the limitation is intrinsic to the Chi.Bio spectrometer sensitivity.

**Figure S5:**
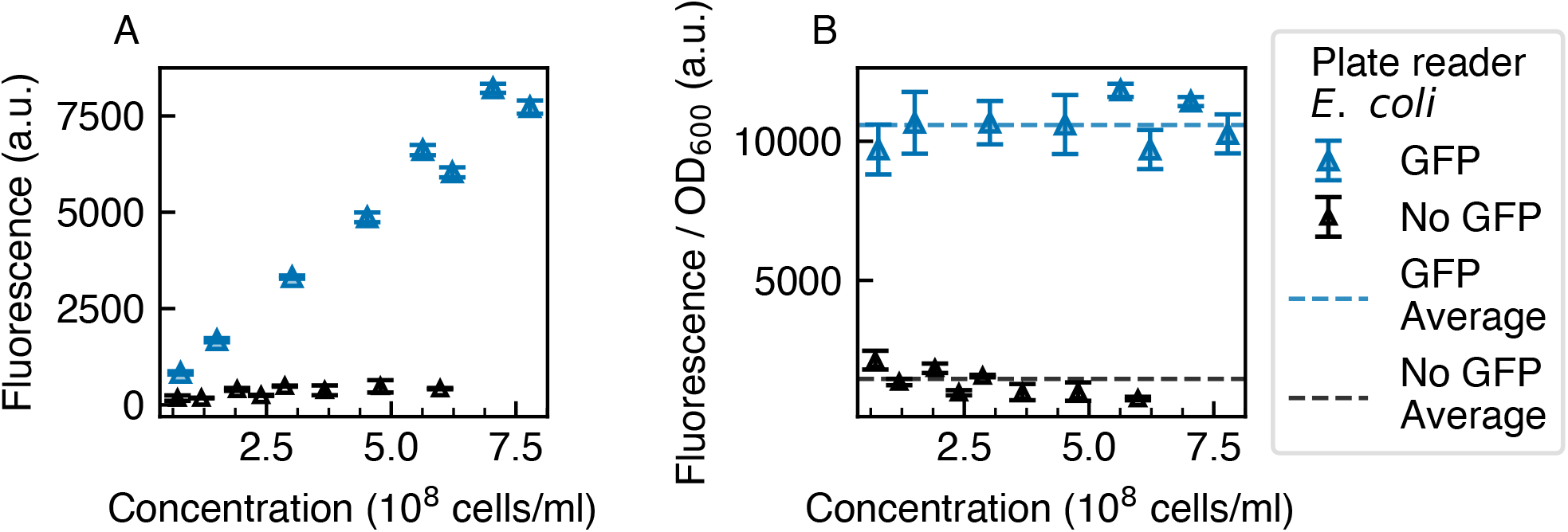
Fluorescence measurements from *E. coli* cells expressing GFP in a plate reader are distinguishable from background. (A) Raw fluorescence intensity of GFP-expressing cells (green) and wild-type controls (black open squares), averaged biological replicate. (B) Same data normalized by OD_600_.

**Figure S6:**
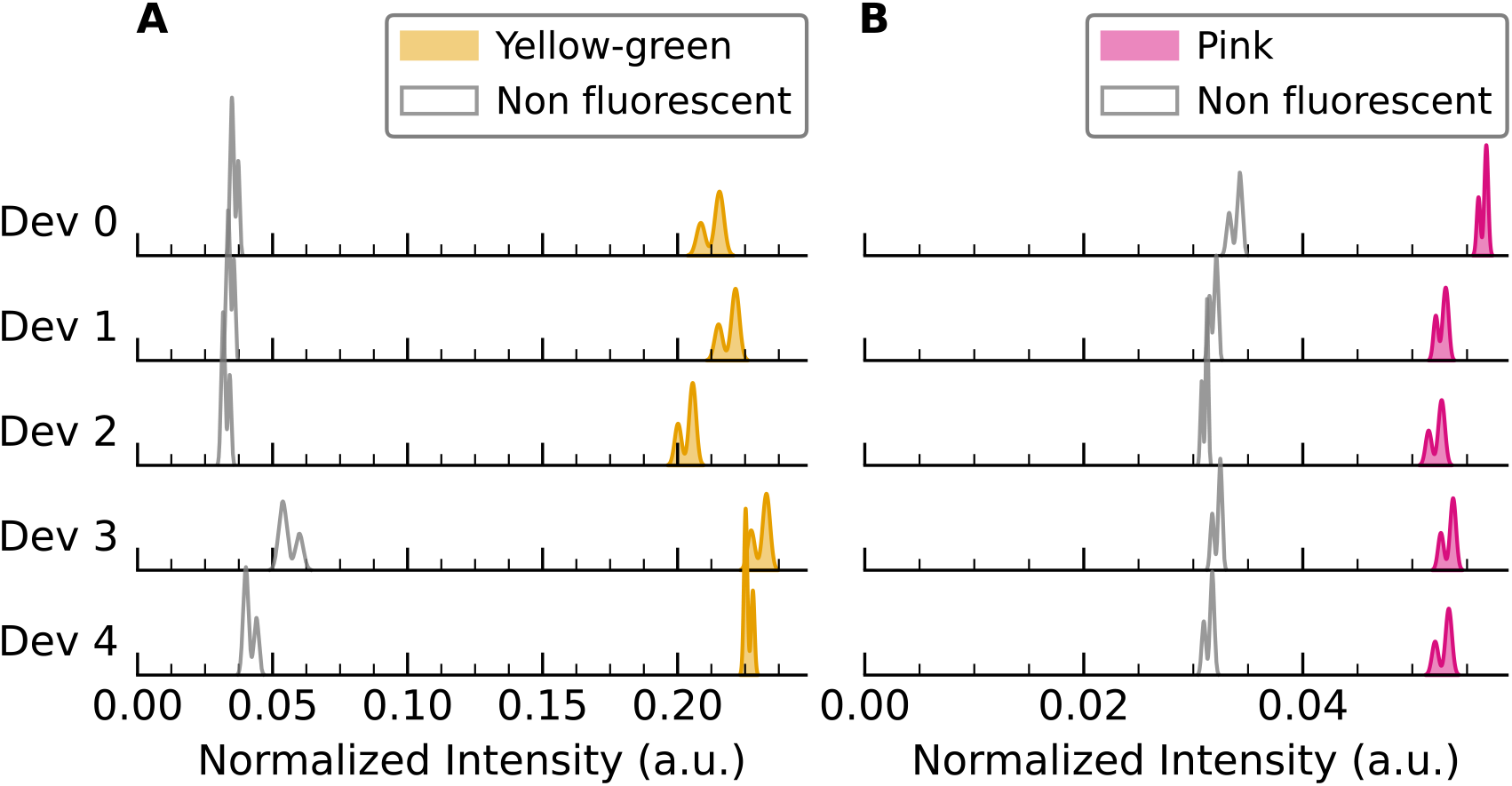
Inter-device variability does not limit the effective separation of fluorescence signal from background in calibration microspheres. (A) Kernel Density Estimation (KDE) of the normalized intensity for yellow-green (filled) and non fluorescent (empty) microspheres, shown for each device. (B) Kernel Density Estimation (KDE) of the normalized intensity for pink (filled) and non fluorescent (empty) microspheres, shown for each device.

## Footnotes

1 H. Steel, personal communication

2 H. Steel, personal communication

